# Engineering orthogonal ribosomes for real-time monitoring using fluorescence

**DOI:** 10.1101/2023.11.19.567736

**Authors:** Eszter Csibra, Bjarne Klopprogge, Georgie Hau Sørensen, Thomas E. Gorochowski

**Affiliations:** Department of Bioengineering, Imperial College Centre for Synthetic Biology, Imperial College London, London, UK; Institute of Biochemistry and Molecular Biology, University of Hamburg, Hamburg, Germany; School of Biological Sciences, University of Bristol, 24 Tyndall Avenue, Bristol, UK; BrisEngBio, School of Chemistry, University of Bristol, Cantock’s Close, Bristol, UK

**Author notes:** Correspondence should be addressed to E.C. and T.E.G. Equal contribution (Bjarne Klopprogge, Georgie Hau Sørensen).

**Keywords:** ribosome, translation, fluorescence, aptamer, biometrology, synthetic biology

## Abstract

A promising route to tackle the trade-off in cellular resources between synthetic protein production and cellular growth is to use a separate dedicated pool of orthogonal ribosomes to produce synthetic proteins. However, the optimisation of strains containing two ribosomal pools – native for the host cell’s proteome and orthogonal for synthetic proteins – has yet to be thoroughly explored. Here, we address this by creating orthogonal ribosomes that fluoresce by inserting fluorescent RNA aptamers into tethered orthogonal ribosomal RNA (TO-rRNA). To study the tolerance of the engineered ribosomes to aptamer insertion, we assembled and screened a library of candidate insertion sites, identifying several sites in both the 16S and 23S TO-rRNA that enables ribosome labelling with minimal effect on translation activity. Serendipitously, we identify one site in 23S TO-rRNA, where insertion appears to not only be tolerated but to enhance orthogonal ribosome activity, across multiple bacterial strains and RNA insertions. Using bulk and single cell assays, we demonstrate that this variant allows us to label orthogonal ribosomes for dynamic tracking and across populations, making it a promising tool for optimising orthogonal translation in engineered cells. Ribosome engineering offers great potential, both for the development of next-generation microbial cell factories, as well as a tool to expand our understanding of ribosome function in living cells. This work provides a window into the assembly, localisation and function of these molecular machines to meet these aims.

## Introduction

Proteins form both key structural elements in cells and power many of the biochemical and biophysical processes essential for life. This central role has led to the control of protein expression, and particularly the translation of protein from messenger RNA (mRNA) by ribosomes, becoming a fundamental part of bioengineering efforts (Green et al. 2014; Mutalik et al. 2013; Salis et al. 2009; Zhao et al. 2022). In this context, one of the most interesting developments over the last decade has been the discovery and optimisation of systems that allow for the generation of a second pool of ribosomes within engineered *Escherichia coli* cells (Liu et al. 2018). Many these systems have been facilitated by two key innovations: orthogonal translation initiation and tethered ribosomal RNA (Fried et al. 2015; Orelle et al. 2015; Rackham & Chin 2005). The tethered orthogonal (TO-) rRNAs contain an orthogonal anti-Shine Dalgarno sequence in the 16S rRNA that allows them to direct the translation of mRNA constructs bearing orthogonal Shine Dalgarno sequences (oSDs, alternatively known as orthogonal ribosome binding sites, oRBSs), while the attachment of the 16S and 23S rRNAs enables 23S rRNA to be mutated without dominant negative phenotypes caused by native and orthogonal subunit mixing.

The field of synthetic biology has seen growing interest in using TO-ribosomes as a basis for engineering cells to contain two ribosome pools: a native pool for performing translation of the host proteome, and an orthogonal pool made up of TO-ribosomes for translating synthetic proteins that may need to be regulated in a different way to endogenous processes (Liu et al. 2018). Using this approach it has been shown that it is possible to decouple the translation of synthetic genes from fluctuations in native ribosome availability and more reliability control gene expression (Darlington et al. 2018). While a promising approach, relatively little is known about the fate of TO-ribosomes in engineered *E. coli* cells, specifically in terms of their production, assembly and localisation. Furthermore, the use of TO-ribosomes is currently constrained by limited protein production yields compared to native translation (Carlson et al. 2019). However, it has been observed that TO-ribosome function may be improved by attention to ribosome assembly and subunit mixing (Aleksashin et al. 2019; Kolber et al. 2021; Schmied et al. 2018), suggesting that knowledge of the cellular fate of engineered rRNAs may be a productive route to enable their optimisation for *in vivo* applications.

Accurate quantification of endogenous and engineered cellular processes, like protein translation, has also seen growing importance in synthetic biology. This stems from the need for measurements of the concentrations and rates of biochemical components to develop predictive models that can better guide bioengineering efforts (Ahn-Horst et al. 2022; Farasat et al. 2014; Muldoon et al. 2021; Nielsen et al. 2016; Schreiber et al. 2016). Without such models, designing large and complex biological systems is near impossible, hampering our ability to tackle important challenges in the field. In this regard, recent work has demonstrated that the careful use of sequencing-based methods (Espah Borujeni et al. 2020; Gorochowski et al. 2017, 2019) can provide a more holistic and quantitative picture of large genetic systems, allowing us to optimise their function more effectively by pinpointing points of failure. Similarly, it has been shown that accurate counts of protein copy numbers in engineered cells can be extracted from bulk fluorescence measurements (Csibra & Stan 2022), and furthermore, that it is in principle possible to infer absolute molecule counts for any fluorophore (Csibra and Stan, manuscript in preparation). Single-molecule approaches have also provided detailed insights into DNA (e.g., plasmid), RNA and protein counts and localisation within living cells (Bienko et al. 2013; Cai et al. 2006; Raj et al. 2008; Raj & van Oudenaarden 2009; Shao et al. 2021). Such developments demonstrate the wide range of tools now at our disposal to study how components like the TO-ribosomes might function and become integrated into engineered cellular systems.

In this work, we develop methods to tag and quantify TO-ribosomes via the attachment of a fluorescent RNA aptamer (**Fig. 1**). While native 5S and 16S rRNAs have been tagged with fluorescent RNA aptamers before (Filonov et al. 2014; Okuda et al. 2017), 23S and TO-rRNAs have not, and we provide the first comprehensive study that aims to identify functional insertion sites throughout a TO-ribosome. To validate the utility of our TO-ribosome variants, we screened them for both the maintenance of ribosomal function by measuring protein translation activity from an oSD reporter, and aptamer folding efficiency by quantifying fluorescence in the presence of the aptamer’s cognate dye. We also assessed the ability for fluorescence signals from our tagged TO-ribosomes to be monitored using plate readers and flow cytometry across *E. coli* strains and growth conditions. Our results provide a step towards the accurate quantification of TO-ribosome abundance in cells and allow for the optimisation of TO-ribosome generation by illuminating its abundance across time (dynamics) and space (localisation), as well as cell populations.

**Figure 1.**
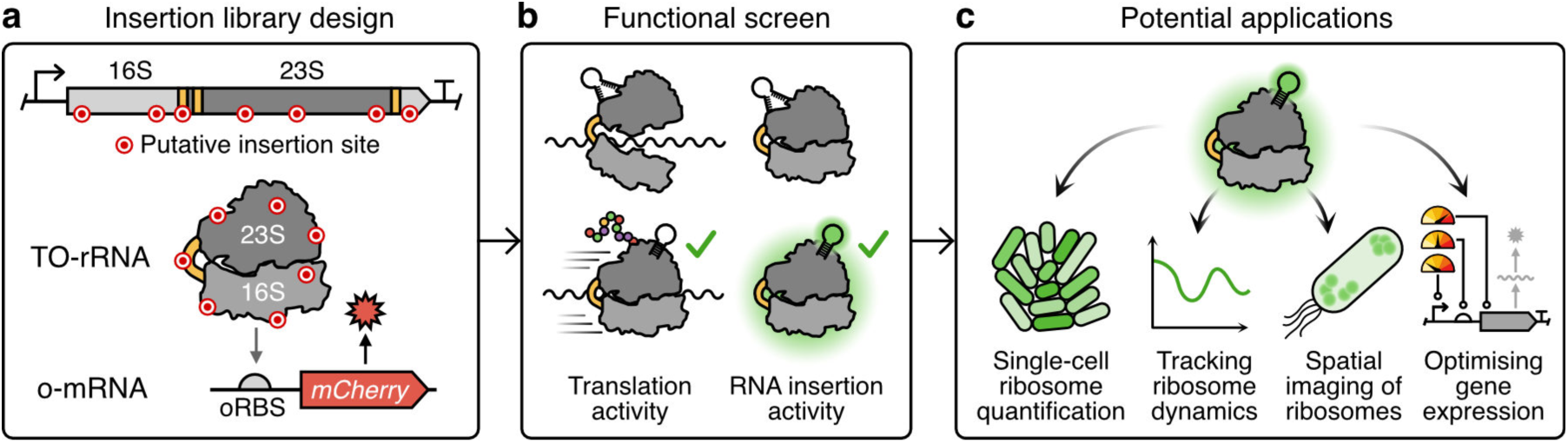
Overview of ribosome functionalisation from candidate site identification to application. (**a**) Identification of candidate sites in the tethered orthogonal (TO-) rRNA that may be permissive for RNA insertion was carried out by structural analysis or sequence analysis. (**b**) Experimental strategy for testing of permissive sites included assays of translation activity and function of inserted Broccoli aptamer. (**c**) The ability to create TO-ribosomes that fluoresce opens up numerous applications from monitoring to optimisation of cellular processes.

## Results

### Designing a TO-ribosome insertion library

To build a fluorescently labelled orthogonal ribosome that can be used to monitor orthogonal translation in real-time in living cells, we first needed to establish which sites in the TO-rRNA are permissive to small RNA insertions. We used two complementary methods to identify candidate sites for this purpose – one based on structural features of the ribosome and the other using sequence information (**Fig. 2**).

**Figure 2.**
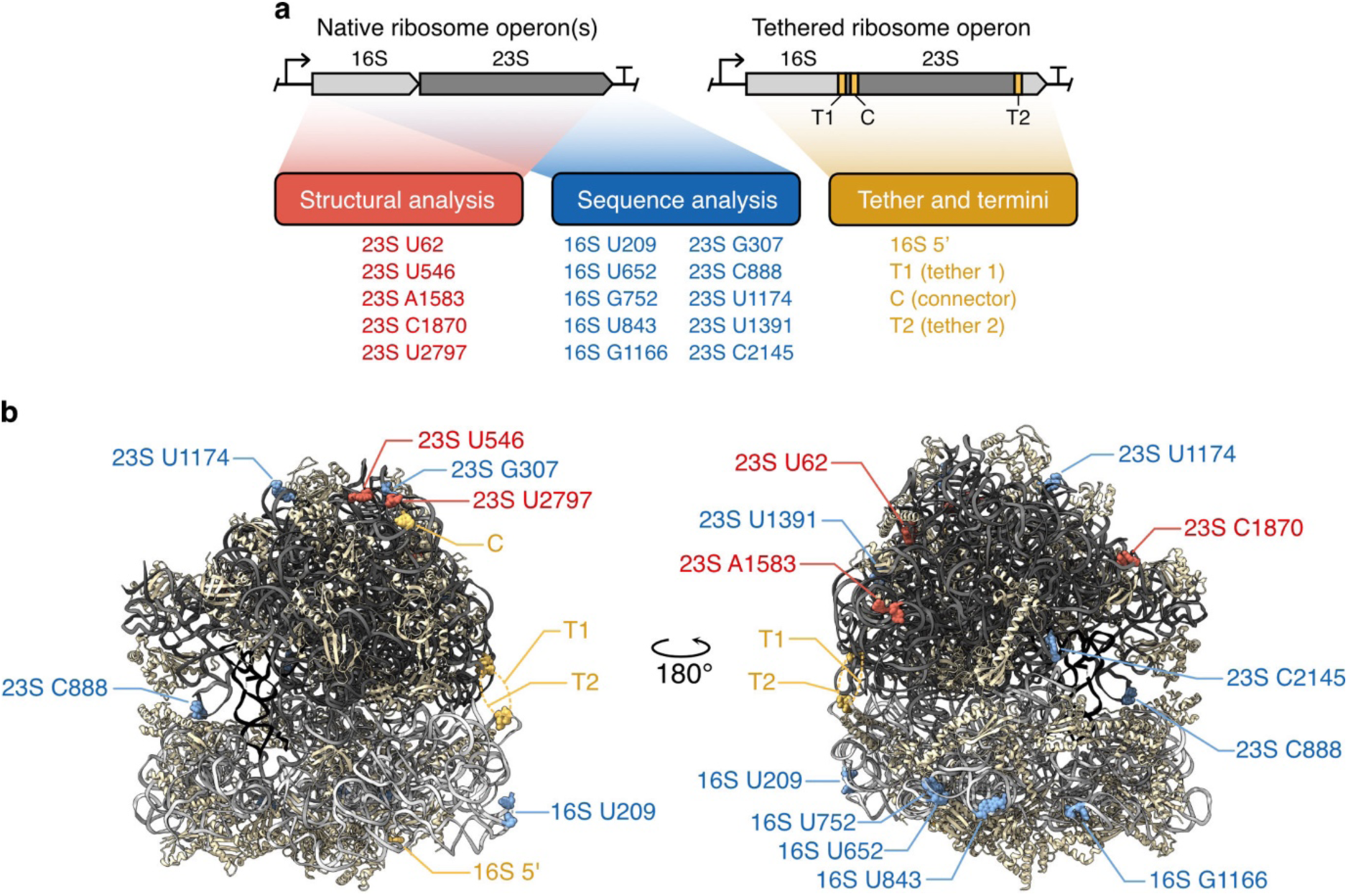
Identification of candidate sites for insertion screening. (**a**) Candidate site identification. Candidate sites were identified by structural (red) and sequence analysis (blue) of the native *E. coli* ribosomal operons. In addition, the 5’ terminus of the TO-ribosome, and all three linker regions (tethers 1 and 2, and the connector; mustard-coloured) were also tested. Operon structures are coloured by ribosomal location: linker region, mustard; 23S rRNA, dark grey; 16S rRNA, light grey. (**b**) Location of identified sites on the 70S tethered ribosome structure (PDB: 8B0X). Tethers T1 and T2 are not modelling in this high-resolution structure and so a dashed mustard-coloured lines denote the points they would attach to in the 23S and 16S subunits. TO-, tethered orthogonal.

Previous efforts to circularly permute rRNA indicated 23 possible insertion sites within the 23S rRNA that may allow translational activity to be maintained (Orelle et al. 2015). We began by assessing the biochemical environment of each of these sites using structural information from cryo-EM structures of native *E. coli* ribosomes (PDB: 4V9D). First, all sites with nucleotides involved in base pairing were discarded. Second, the distance to the closest residue, which is not part of the same loop/helix, was measured. Third, the overall position within the ribosomal structure was assessed. In total, five of the 23 sites (23S U62, U546, A1583, C1870 and U2797) located in superficial loops were selected for testing (**Fig. 2a**, left hand panel).

We also performed a multiple sequence alignment of 23S and 16S rRNAs from *Escherichia* genus isolates using the SILVA rRNA database (Quast et al. 2013) to find permissive sites. These would be highlighted by the existence of natural insertions present in organisms related to our model *E. coli* system. Sites where frequent insertions were observed, as compared to the consensus 16S and 23S sequences, were selected for further testing – constituting a further ten sites (**Fig. 2a**, middle panel).

Finally, four additional sites were selected based on the design of the oRiboT2 tethered ribosome (Orelle et al. 2015): an insertion site at the 5’ terminus (5’), and in each of the linker regions, i.e., in the T1 and T2 tethers and the connector (**Fig. 2a**, right hand panel). In total, our designed library contained 19 candidate insertion sites, constituting sites covering both subunits (**Fig. 2b**).

### Assessing sites at which ribosomes may be functionalised

We built our library by inserting a Broccoli RNA aptamer sequence (Filonov et al. 2014) into the oRiboT2 tethered orthogonal (TO-) ribosome (Orelle et al. 2015), separately at each of the 19 candidate sites. To test the activity of the resultant TO-ribosome variants, cells were co-transformed with an orthogonal mCherry reporter construct (containing an orthogonal SD) to assess the performance of the variants compared to the ‘wild-type’ (WT) parental TO-ribosome (**Fig. 3a**). Most of the candidate sites retained over half of their activity, and we observed a strong correlation between the two tested strains, DH10B and BL21(DE3) (**Fig. 3b** and **Supplementary Fig. 1**). We observed a higher rate of permissiveness from sites in the 23S rRNA segments of the TO-rRNA, as opposed to the 16S segments. However, we would need to expand the library size to investigate whether this is a general trend. All of the linker regions supported translation activity, with the connector site, C, that was used to attach the 5’ and 3’ ends of the 23S rRNA for permutation (Orelle et al. 2015), performing better than the tethered regions. We later found that two of the sites that performed well (23S U1174 and C2145) had been previously utilised for the insertion of protein-binding tags for ribosome purification (Matadeen et al. 2001 p. 2008; Yokoyama & Suzuki 2008), showing we could recapitulate positive results previously obtained by others. A remaining set of 6 insertion sites (16S 5’, U209, U652, and 23S U546, C888, A1583, C1870) that we can identify as retaining high levels of translation activity (that is >80% in BL21(DE3) and >65% in DH10B) have not, to our knowledge, been previously tested as candidate sites for insertion tolerance, and we therefore identify for the first time as effective sites for ribosome engineering.

**Figure 3.**
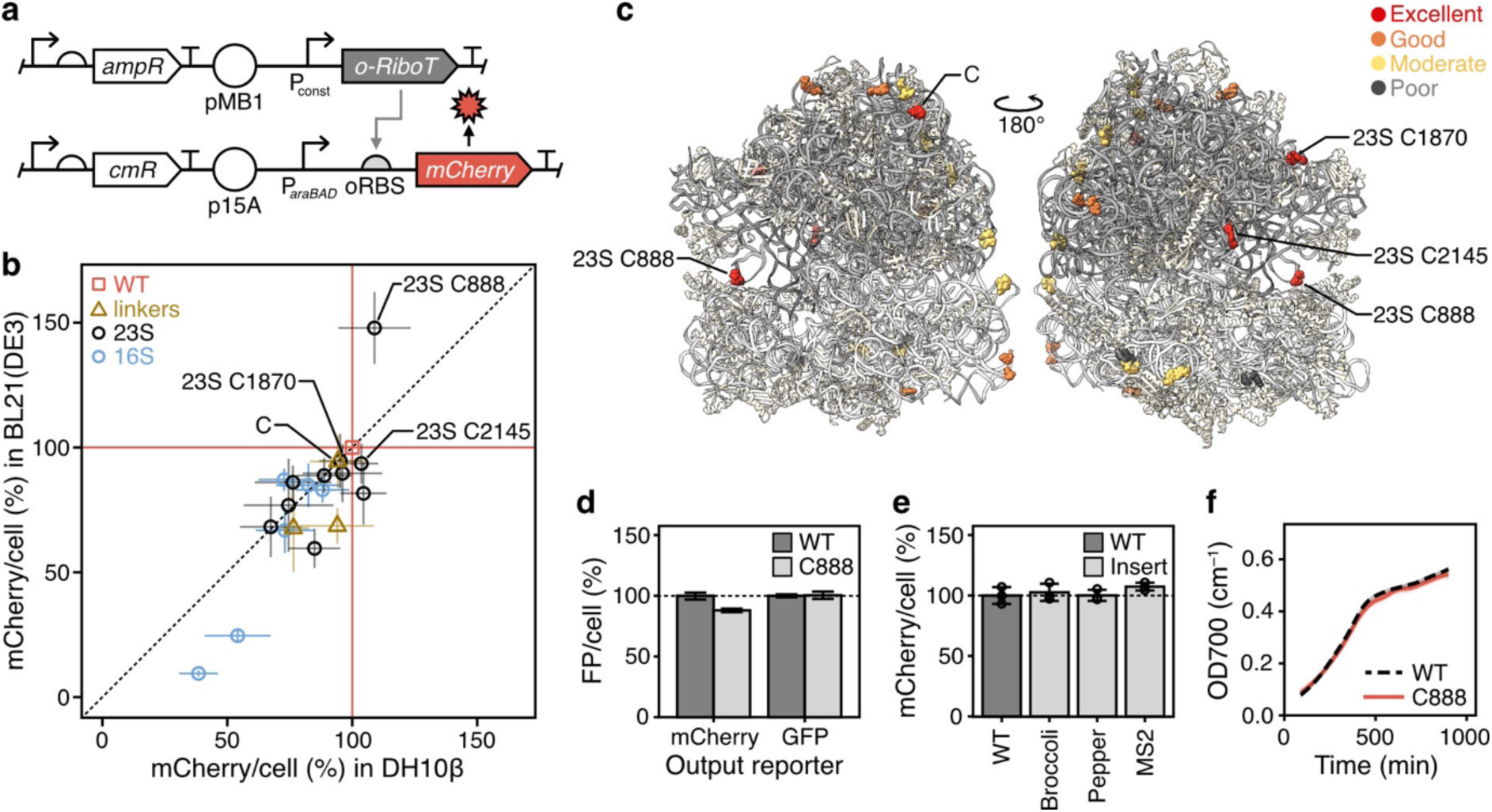
Insertion screening of tethered orthogonal rRNA reveals permissive sites at which ribosomes may be functionalised. (**a**) Schematics of the genetic constructs used to express the TO-ribosomes from the *o-RiboT* gene (top) and test their ability to express an oSD RBS (oRBS) driven mCherry reporter (bottom). (**b**) Performance of mutant TO-ribosomes in two *E. coli* strains. Cells were co-transformed with orthogonal ribosome and orthogonal mCherry reporter plasmids, and grown in M9 media, while oSD-mCherry was induced with 0.1% arabinose. mCherry production per cell was calculated from mCherry and OD readings, and results were compared to the readings from the ‘WT’ TO-ribosome variant at 720 min, which was set to 100%. Plotted data represents the mean and standard deviation of results over at least 2 independent repeats conducted in quadruplicate. Sites are coloured by ribosomal location: no insert/WT, red; insert in linker region, yellow; insert in 23S rRNA region, light blue; insert in 16S rRNA region, dark blue. (**c**) Structure of TO-ribosome, showing insertion sites by performance (red to dark grey, best to worst; PDB: 8B0X). Variants were classified as Excellent (>90% activity in BL21(DE3) and >85% activity in DH10B strains), Good (>80% activity in both strains), Moderate (>60% in BL21(DE3) and >70% in DH10B) and Poor (<60% in both). (**d**) Performance of WT and C888 insertion mutant when expressing mCherry and GFP orthogonal reporters in DH10B strain. Cells were grown and induced, and data was analysed, as in panel (b). (**e**) Permissiveness of C888 site for a range of insertions in DH10B strain. Cells were grown and induced, and data was analysed, as in panel (a). (**f**) Growth curve of WT and C888::Broccoli variants in DH10B strain over a standard assay. Representative of multiple experiments. WT, poRiboT2; C888, poRiboT2_C888::insertion. Insertions in panels (b)–(d) and f are Broccoli. In panel (d), the identity of the insertion is given on the *x*-axis.

Curiously, another site used previously for ribosome isolation (23S U2797, in the apex loop of 23S helix H98) (Ali et al. 2006; Matadeen et al. 2001; Shi et al. 2012; Youngman et al. 2004; Youngman & Green 2005), was not identified as one of our highest performing variants. Because few of these previous studies conducted comparative analyses similar to ours, it remains possible that this site is not one of the best sites for isolation, given a comprehensive screen. Nonetheless, it remains interesting that other groups have identified this site as functional for the purposes of ribosome activity assays in cell free assays, given its position in its ranking as one of the least effective of our tested sites in DH10B cells (**Supplementary Fig. 1b**). This suggests that most of our tested sites may be useful for studies in which a small degree of reduced functionality may be tolerated. Alternatively, there may be differences between the functional implications of inserted RNA sequences *in vitro* and *in vivo*, or between native and tethered ribosomes, that have yet to be explored.

### Exploration of the translation enhancing 23S C888 site

Insertion of the Broccoli aptamer at the 23S C888 site revealed the most striking data from our library. The 23S C888 site performed strongly in both strains and reliably exceeding the translation activity of the parental WT ribosome in BL21(DE3) cells. For this reason, it was selected for further testing. All subsequent experiments were carried out in DH10B cells using our pSEVA361 based reporters. This reporter system has the advantage of having been thoroughly characterised from our earlier work, and is part of a library of fluorescent protein reporters (Csibra & Stan 2022).

First, we checked whether the high activity of the C888 variant was maintained across multiple reporters by testing its activity on an alternative fluorescent protein reporter construct containing an upstream orthogonal SD site and normalised its result to the parental TO-ribosome (**Fig. 3c**). This showed that the C888 variant is functional across different reporters, with 88% activity for mCherry and 101% for mGFPmut3. Next, we asked whether TO-ribosomes were tolerant to insertions of other RNA sequences at the C888 position. To test this, we assembled variants containing the Pepper aptamer (Chen et al. 2019) and MS2 binding site (Witherell et al. 1991) and measured the TO-ribosome activity on our mCherry reporter construct. We found that for all inserts, the TO-ribosome showed excellent activity that matched or even exceeded the non-modified TO-ribosome (**Fig. 3d**). None of our tests showed significant impacts on growth rate (**Fig. 3e**), ruling out reduced dilution due to cellular replication as the source of the high protein per cell values. These results suggest that the C888 position is amenable to a range of RNA insertions with diverse sequences and structures between 43–71 nt long.

### The Broccoli RNA aptamer is functional within the rRNA scaffold

Having established that TO-rRNA retains translation activity in the presence of inserted sequences at position C888, we next asked whether the inserted RNA sequences fold and function correctly in the context of the TO-rRNA. Fluorescence from the Broccoli RNA aptamer is reportedly enhanced by placing it in the context of a structured scaffold such as F30 (Filonov et al. 2015). However, we reasoned that this should not be necessary in the context of a stable structured RNA such as a TO-rRNA.

We tested the functionality of the aptamer by growing cells containing our TO-ribosomes in the presence of DFHBI-1T (**Fig. 4a**) and monitored green fluorescence alongside bacterial growth over time in a plate reader. Two constructs were tested: (i) a constitutive expression construct, based on the original poRiboT2 construct (**Methods**) in DH10B cells, and (ii) an inducible construct in which the promoter had been replaced with an IPTG-inducible P*tac* promoter in DH10B Marionette cells where LacI is expressed from the genome (Meyer et al. 2019). Broccoli levels per cell were estimated by calculating normalised green fluorescence over normalised OD700 values, and removing the signal from a paired negative control, identical to the tested sample with the exception of the Broccoli insertion (**Fig. 4b**). We found a clear increase in fluorescence signal per cell from cells with Broccoli-functionalised ribosomes. They also showed expression dynamics consistent with constitutive (**Fig. 4b**; left) and inducible (**Fig. 4b**; middle and right) systems. This demonstrates that Broccoli is functional within the TO-rRNA scaffold, without the addition of an F30 scaffold, and that the insertion of a single Broccoli (rather than a tandem or larger arrays of aptamers) may, in some cases, be sufficient to study and optimise TO-ribosome expression in living cells.

**Figure 4.**
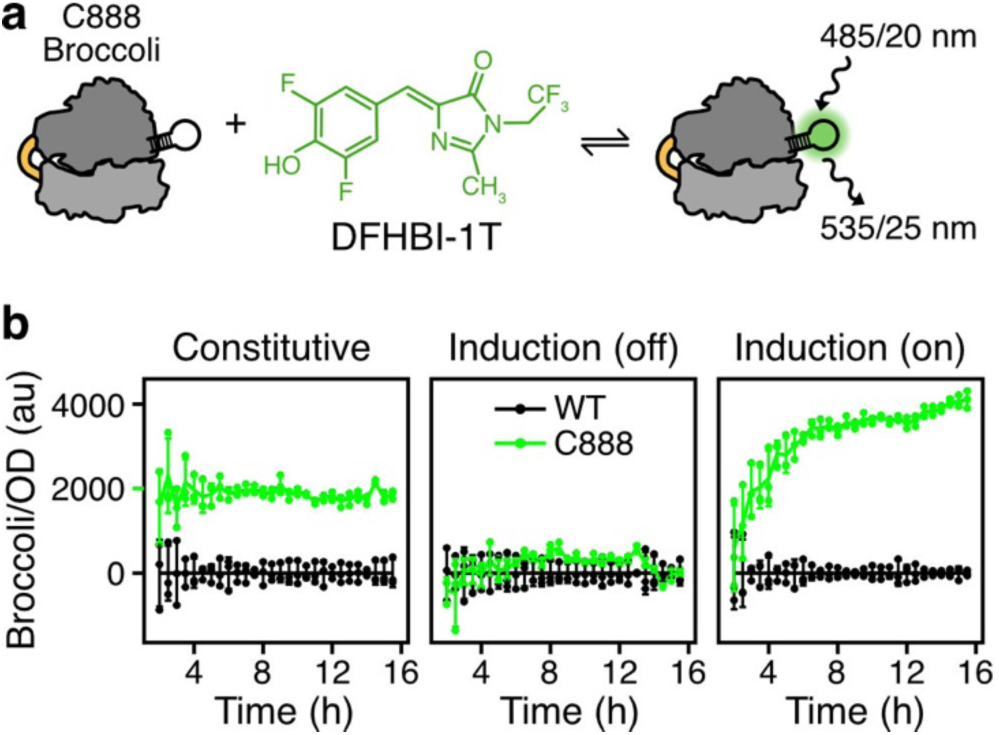
O-Ribosomes can be detected in live cells with Broccoli. (**a**) Diagram of Broccoli fluorescence with DFHBI-1T binding. (**b**) Broccoli fluorescence can be detected from cells expressing TO-ribosomes with C888::Broccoli insertion, from both a constitutive promoter (phage lambda pL promoter, without repressor) or an IPTG-induced P*tac* promoter. Cells (DH10B, left panel; DH10B-Marionette, middle and right panel) were transformed with orthogonal ribosome variants (poRiboT2: left panel, or pTac_oRiboT2, middle and right panel, with or without C888::Broccoli insertions), and grown in M9. Orthogonal ribosome expression was either not induced (left, middle panels) or induced with 1mM IPTG (right panel). Broccoli per cell was quantified by normalising green fluorescence per OD700 readings from Broccoli containing samples to matched controls. Plotted data represents the mean, standard deviation, and triplicate data points over time. WT, oRiboT2 construct without Broccoli insertion; C888, constructs containing C888::Broccoli.

### Monitoring TO-ribosome concentration dynamics

Having established that Broccoli labelling allowed us to track labelled ribosomes, we next asked what Broccoli fluorescence could tell us about the abundance of orthogonal ribosomes during constitutive expression across a range of growth conditions in widely used *E. coli* strains.

We observed striking differences in ribosome abundance across strain, sugar and growth phase (**Fig. 5a**). In DH10B cells, constitutive expression resulted in an approximately 2-fold higher level of TO-ribosome abundance per cell in fructose than in glucose-supplemented media. This suggests that carbon source can affect the production and/or degradation of TO-ribosomes. In contrast, this pattern was not seen for the BL21(DE3) strain, with similar TO-ribosome concentrations across the carbon sources. Instead, we observed a prominent effect of growth phase, from log phase levels that are comparable to those of the DH10B strain grown in fructose, followed by a drop to near zero during stationary phase (**Fig. 5a**), a response that was reproducible over multiple experiments.

**Figure 5.**
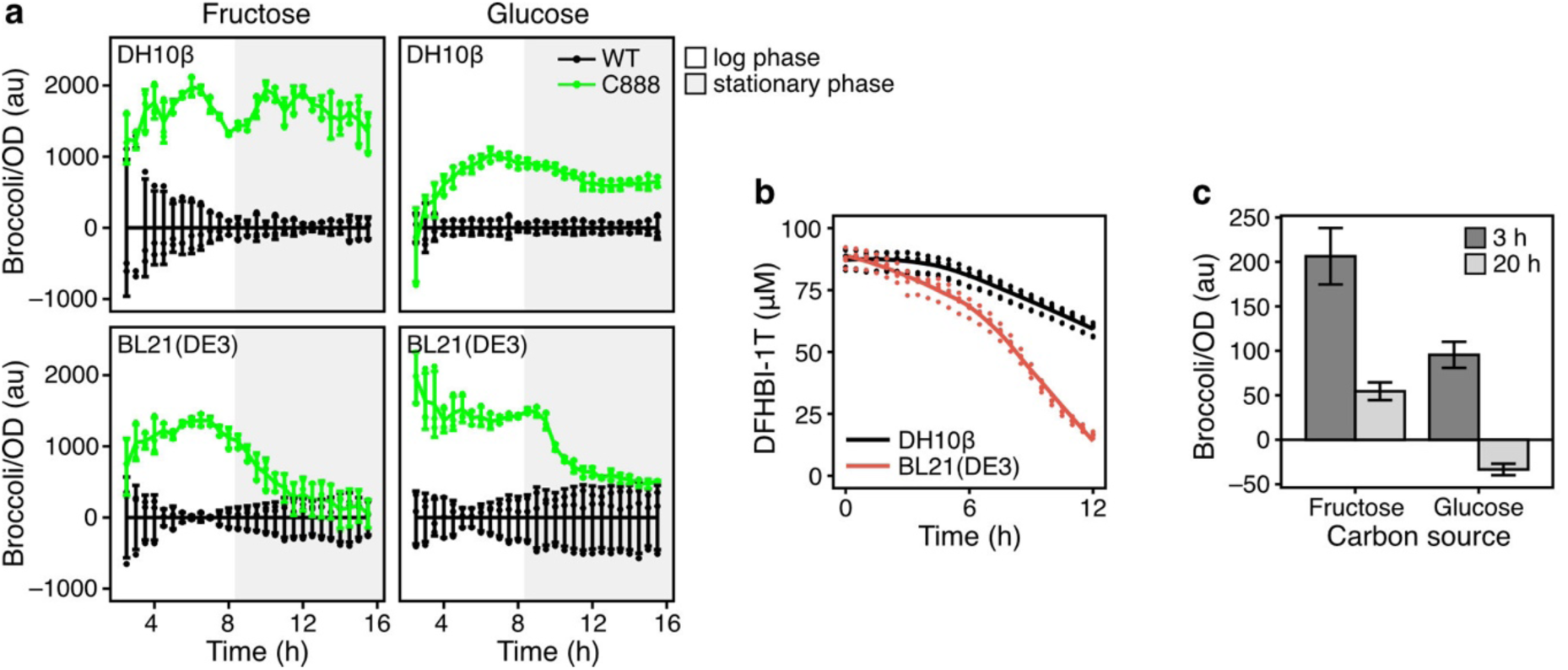
Application of Broccoli insertion to orthogonal ribosome quantification in engineered cells. (**a**) Time course tracking of Broccoli fluorescence under a range of growth conditions suggests TO-ribosome abundance may be affected by bacterial strain, sugar and growth phase. DH10B cells were transformed with orthogonal ribosome variants (poRiboT2 with or without C888::Broccoli insertion), and grown in M9 supplemented with 0.8% fructose or glucose as a carbon source. Broccoli per cell was quantified by normalising green fluorescence per OD700 readings from Broccoli containing samples to matched controls. Plotted data represents the mean, standard deviation, and triplicate data points over time. Data is representative of at least three independent experiments. (**b**) DFHBI-1T tracking over extended time course assays in plate readers. OD426 was monitored over time, and DFHBI-1T concentration was calculated by normalising to media OD426, subtracting the cellular OD426 contribution, and converting the resultant DFHBI-1T OD426 from absorbance to concentration via its extinction coefficient (see **Methods**). (**c**) Flow cytometric analysis of ribosome abundance. DH10B cells were grown as in panel (a) and aliquots were removed at 3 h (log phase) and 20 h (stationary phase) for flow cytometric analysis. DFHBI-1T was added to all samples (200 µM) and green fluorescence was quantified for cells with and without Broccoli, the latter used to normalise fluorescence for the former. Data represents the mean and standard deviation of a triplicate dataset. WT, poRiboT2; C888, poRiboT2_C888::Broccoli.

During these experiments, we observed that the bright yellow colour of media containing DFHBI-1T was preserved after overnight growth in wells containing only media but appeared diminished in the presence of bacterial cultures. If DFHBI-1T was degrading over an extended time course assay, this may partly explain the decrease in Broccoli signals we observed in the BL21(DE3) cultures. Using a method adapted from the absorbance-based fluorescent protein quantification assay (Csibra & Stan 2022) (see **Methods**), we tracked DFHBI-1T levels across DH10B and BL21(DE3) cultures at a range of starting concentrations (**Fig. 5b** and **Supplementary Fig. 2**). While it is difficult to track high concentrations of DFHBI-1T due to the sum of the absorbance from the cell and the label being too high to quantify and leading to missing values in the data (**Supplementary Fig. 2**, 200µM), it is possible to track lower concentrations accurately. It is clear from this data that DFHBI-1T concentrations drop over time for both strains, but that the drop is more pronounced for BL21(DE3). Nonetheless, we observe that DFHBI-1T is stable in DH10B cultures during the first 6 hours of log phase growth allowing for extended monitoring of TO-ribosome concentrations over time.

To investigate ribosome abundance using a method that does not require continuous incubation of DFHBI-1T over several hours, we turned to flow cytometry to validate our plate reader results. Using DH10B cells grown in fructose-supplemented M9, cultures were grown identically to plate reader cultures, but without the addition of DFHBI-1T, and aliquots were removed at two timepoints for flow cytometric analysis with DFHBI-1T. The flow cytometry data recapitulates the approximately 2-fold decrease in ribosome abundance between fructose- and glucose-supplemented cultures (**Fig. 5c**, supporting the notion that carbon source affects orthogonal ribosome abundance through either production and/or degradation rate differences in DH10B cells.

### Optimising reporters for measurement of orthogonal ribosome activity

Throughout these experiments, we observed varying background expression levels of our oSD reporters by native ribosomes in strains not harbouring orthogonal ribosome vectors. In certain cases, this reached significant levels of up to ∼50% of total reporter expression, compared to cells with orthogonal ribosomes present (**Supplementary Fig. 3b**, mCherry reporter). It has been shown that the first internal methionine codon in the standard mCherry sequence serves as a site of internal translation initiation and that replacing the AUG for a CUG (leucine) codon reduces the internal initiation frequency to less than 10% of that in the standard mCherry (Fages-Lartaud et al. 2022). We reasoned that an mCherry_M10L construct would be useful to assess whether our data with the standard mCherry was the result of the binding of native ribosomes to (i) our orthogonal SD upstream of the canonical AUG codon, or (ii) the internal AUG that happens to be preceded by an AG-rich sequence (GAGGAAGATAAC) coding for the previous four codons (EEDN). In the case of (i) we would expect the M10L construct not to affect the mCherry expression patterns, while in the case of (ii), we would expect a reduction of mCherry expression for all conditions. Experiments with the new construct (**Supplementary Fig. 3b**, mCherry_M10L reporter) supported the existence of an internal SD upstream of M10. Removal of this initiation site increased our signal to noise ratio for mCherry detection by 3.2-fold.

## Discussion

In this work, we have shown that TO-ribosomes are functionally robust to many different insertions into their TO-rRNA. We demonstrate that selection of permissible sites using either structural or sequence information typically leads to <20% decrease in translation activity. Our top performing site, C888, is located at the apex loop of 23S helix H38, also known as the A-site finger, and forms part of B1a, one of the twelve ribosomal inter-subunit bridges. This structure is reported to be involved in ribosome assembly (Yassin & Mankin 2007), subunit association (Liiv & O’Connor 2006; Sergiev et al. 2005) and reading frame maintenance (Komoda et al. 2006). Although H38 truncation appears to be have modest phenotypic effects on translational activity (Komoda et al. 2006; Liiv & O’Connor 2006; Sergiev et al. 2005), it has been observed to impair ribosome assembly (Yassin & Mankin 2007), and curiously, has also been observed to increase translocation rates of mutant ribosomes *in vitro* (Komoda et al. 2006; Kudrin et al. 2018; Wang et al. 2011). We find that extensions of this loop lead to notable increases in translation activity that are strain dependent (∼50% in BL21(DE3) and ∼6% in DH10B). As far as we are aware, the phenotypic effects of H38 extensions have not previously been investigated and could result from either an effect on translocation rate or ribosome assembly.

By using a Broccoli aptamer as a common insert in our library, we are also able to show that TO-ribosome concentrations can be monitored via fluorescence in both plate readers and by flow cytometry. Broccoli insertion into ribosomal targets has been previously reported, with the first document approach targeting the 5S rRNA (Filonov et al. 2014) and another exploring the robustness of the ribosome to different types of aptamers at a single location (Okuda et al. 2017). Our results are the most comprehensive to date, elucidating many new sites for future TO-ribosome engineering that appear to have little impact on translation activity and which are scattered across both major subunits.

An unexpected outcome of this work was the impact that different strains and media had on TO-ribosome concentrations and dynamics during batch cultures grown to stationary phase. Cells grown in glucose showed lower accumulation of TO-ribosomes than those grown in fructose. There is a large body of literature that connects native ribosome abundance with growth rate and growth conditions (Dai et al. 2016; Kim et al. 2020; Scott et al. 2010; Weiße et al. 2015). It has been proposed that during high growth rates, native ribosomes spend proportionally more time producing more ribosomes (ribosomal proteins) than other proteins, and this ratio decreases with decreasing growth rate (Scott et al. 2010). Initially, our fructose data seems to support this, as levels decrease with increasing growth rates. However, we are quantifying rRNA production, not protein production, and increased ribosomal protein production might be expected to result in higher levels of orthogonal ribosomes too. An alternative hypothesis might be that orthogonal ribosome abundance is dependent on its substrate concentration: ribosome abundance has been linked to ribosome demand, as inactive ribosomes are more prone to degradation (Zundel et al. 2009). As our reporter expression is driven by the *araBAD* promoter, it is inhibited by the presence of glucose via catabolite repression (Lichenstein et al. 1987), leading to low mRNA levels in the presence of glucose, that is supported by the low levels of mCherry measured from cultures grown in glucose (data not shown). Further work will be required to disentangle these effects.

Broccoli and related RNA aptamers have been used successfully over the last decade to monitor RNA localisation in living cells. However, their use in molecule quantification has lagged. We suspect this is largely due to the fact that most interest in this area concerns mRNA quantification, which suffers the dual challenge of low mRNA copies per transcript per cell (<10 in bacteria, (Bremer & Dennis 2008; So et al. 2011; Xie et al. 2008)), and poor folding efficiency of the Broccoli aptamer in the relatively unstructured context of mRNAs (Filonov et al. 2015). In contrast, ribosomal RNA is maintained at far higher copies per cell (in the order of 10^4^) and forms an inherently structured scaffold to enable Broccoli folding, making it a more plausible target for accurate quantitative monitoring.

During this work, we identified a strain-dependent drop in DFHBI-1T levels over time (**Fig. 5b**), which makes continuous monitoring over extended time periods a challenge. It is clear from our data that a drop in DFHBI-1T is found for both strains we tested but is more pronounced for the BL21(DE3) cells. As far as we are aware, this is the first time such an effect has been described in the literature. It is currently not clear why a larger drop occurs for BL21(DE3). However, it may be due to the strains higher growth rate compared to the DH10B cells we also tested, and the ability for the cells to grow to higher final concentrations. While this constitutes a challenge to quantification, it is of interest that DFHBI-1T levels may be effectively tracked in live cultures. This allows us to make informed decisions about which timepoints can be reliably compared and opens the possibility of correcting fluorescence readout for DFHBI-1T concentration at any given time.

As synthetic biology transitions from tinkering with biology to engineering it, the development of quantitative approaches for the real-time monitoring of core cellular components and machinery, like ribosomes, will be key to supporting informed optimisation of complex biomolecular systems in living cells (Shao et al. 2021). This work provides a step in this direction, offering new avenues to understand and tune protein synthesis, and further explore how orthogonal cellular machinery can be best used to create robust and predictable biotechnologies.

## Materials and Methods

### Strains and media

For all plasmid cloning and propagation, *Escherichia coli* strain DH10B (Δ(ara-leu) 7697 araD139 fhuA ΔlacX74 galK16 galE15 e14-phi80dlacZΔM15 recA1 relA1 endA1 nupG rpsL (StrR) rph spoT1 Δ(mrr-hsdRMS-mcrBC) (New England Biolabs, C3019I) was used. For the characterisation of ribosomal variants, *E. coli* strains DH10B or BL21(DE3) (fhuA2 [lon] ompT gal (lambda DE3) [dcm] ΔhsdSlambda DE3 = lambda sBamHIo ΔEcoRI-B int::(lacI::PlacUV5::T7 gene1) i21 Δnin5) (New England Biolabs, C2527I) were used. The expression of pTac induced ribosomes was carried out in DH10B-Marionette strains (Meyer et al. 2019), a gift from Christopher Voigt (Marionette-Clo, Addgene #108251).

All cells were grown in either LB media (Sigma–Aldrich, L3522) for outgrowth and propagation, or M9 minimal media supplemented with fructose or glucose (6.78 g/L Na2HPO4, 3 g/L KH2PO4, 1 g/L NH4Cl, 0.5 g/L NaCl (Sigma–Aldrich, M6030), 0.34 g/L thiamine hydrochloride (Sigma T4625), 0.8% D-glucose (Sigma–Aldrich, G7528) or 0.8% fructose (F3510), 0.2% casamino acids (Acros, AC61204-5000), 2 mM MgSO_4_ (Acros, 213115000), and 0.1mM CaCl_2_ (Sigma–Aldrich, C8106)) for characterisation experiments. Inducers used included L-(+)-arabinose (Sigma-Aldrich, A3256) or isopropyl beta-D1-thiogalactopyranoside (IPTG) (Sigma–Aldrich, I6758). For antibiotic selection, 100 µg/mL ampicillin (Sigma–Aldrich, A9518), 50 µg/mL kanamycin (Sigma–Aldrich, K1637) or 10 µg/mL gentamicin (Sigma– Aldrich, G3632) were used.

### Creation of an orthogonal ribosome insertion library

All cloning steps were performed according to the manufacturers protocol unless specified otherwise. The 71 bp long insertion sequence containing the Broccoli aptamer, flanked by BsmBI restriction sites, was generated by annealing two reverse-complementary primers (all primers were synthesized by Integrated DNA Technology). Equimolar amounts were mixed, heated to 95°C and the temperature was decreased to 50°C (30 s per 1°C). This insertion sequence and the poRibo-T2 plasmid (a gift from Michael Jewett; Addgene plasmid #69347; (Orelle et al. 2015)) were used as PCR templates. PCR was performed using Q5 High-Fidelity DNA Polymerase (New England Biolabs, M0491S) and primers harbouring overhangs for Gibson Assembly. After gel extraction using the Monarch DNA Gel Extraction Kit (New England Biolabs, T1020S) Gibson Assembly (New England Biolabs, E2611S) of the two parts was performed, and 2 µL were used for transformation. After overnight growth at 37°C, individual colonies were picked and a colony PCR using Quick-Load Taq 2X Master Mix (New England Biolabs, M0271L) was used to confirm the expected insert size. Positive colonies were incubated overnight at 37°C and plasmids were isolated using the Monarch Plasmid Miniprep Kit (New England Biolabs, T1010L). For all assembled plasmids the sequence was confirmed by Sanger sequencing. Insertion of other sequences were performed in a similar way. Plasmid sequences are provided in **Supplementary Data 1**.

### Assembly of reporter plasmids

Most reporter plasmids were derived from the SEVA based mCherry reporter plasmids first reported in (Csibra & Stan 2022). The parental vector (pS361_ara_mCherry) is a medium copy p15A vector (pSEVA361) containing a native SD (AGGAGG) followed by an N-terminally His-tagged, codon-optimised mCherry coding sequence (FPbase ID: ZERB6, (Lambert 2019)). The SD of this reporter was swapped to an orthogonal SD (ACCACA) that matches the antiSD within poRIboT2, by reverse PCR and blunt end ligation with KLD (NEB M0554S). Subsequently, an internal initiation site within the mCherry coding sequence was removed by changing the Met10 codon (AUG) to a leucine codon (CUG) to create the M10L variant, using the same protocol. The mGFPmut3 oSD reporter was assembled from the parental vector (pS361_ara_mGFPmut3) similarly, by swapping the native SD for an oSD. For the BL21(DE3) screen, mCherry reporters with an oSD and under the control of a T7 promoter were assembled in a pSEVA661 (p15A, gentamicin resistance) backbone. All plasmid sequences were verified by Sanger sequencing.

### Monitoring cell growth and fluorescence using plate reader assays

Single colonies were used to inoculate 1 mL M9 media containing antibiotics and incubated in deep well plates at 30°C and 700 rpm for 16 h in a shaking incubator. Starter cultures were created by diluting overnight cultures to an OD600 of 0.05/ml in 1 mL fresh M9 media with antibiotics and incubated at 30°C and 700 rpm for an hour in the same shaking incubator, before transfer to a 96-well black polystyrene clear-bottom plate (Corning Inc.) for plate reader assays. Inducers such as arabinose (to 0.1%) and labels such as DFHBI-1T (to 200 µM, Bio-Techne Ltd, 5610) were added to these plates, with the cultures added to achieve final volumes of 200 µL/well.

Plate reader measurements were taken using a multiwell plate reader at 30°C for 16 h with double orbital shaking. The plate readers used for the experiments in this manuscript include a SpectraMax iD5 (Molecular Devices, LLC.), a Synergy Neo2 (Biotek) and a Tecan Spark multimode plate reader (Tecan). Cell density measurements were monitored using OD600 and OD700 measurements. Protein and RNA fluorescence was monitored using green (ex 485/20 nm, em 535/25 nm) and red (ex 560/20 nm, em 610/20 nm) filter sets, with minor variations depending on the instrument used. Plate reader measurements were calibrated for red fluorescence using mCherry lysates, and OD with microspheres, as previously described (Csibra & Stan 2022).

### Monitoring fluorescence by flow cytometry

Samples (2–5 µl) were removed from cultures growing in a plate reader at log phase (3 h post transfer to multiwell plates) or stationary phase (20 h) and diluted in 1 ml M9 with vigorous vortexing. Aliquots from this dilution were transferred to 96-well round-bottom plates containing 4 µl 10mM DFHBI-1T (Bio-Techne) for a final concentration of 200 µM. Fluorescence was analysed on an Attune NxT flow cytometer in the BL1 (ex 488, em 535/25 nm) and YL2 (ex 560, em 620/20 nm) channels.

### Quantification of DFHBI-1T over time in plate reader assays

DFHBI-1T is a small molecule fluorophore whose quantum yield is approximately 100-fold lower in solution than in its Broccoli-bound form (Filonov et al. 2014). However, its light absorbance is efficient even alone in solution, and its absorbance spectrum has been previously recorded as having a peak at 426 nm and an extinction coefficient of 35,400 M^−1^cm^−1^ (**Supplementary Fig. 2a**, data from (Filonov et al. 2014)). In the development of the FPCountR method for fluorescent protein quantification with plate readers (Csibra & Stan 2022), a method for the accurate quantification of fluorescent proteins was developed based on light absorbance at the proteins’ absorbance maxima. It was established that protein abundance may be reliably monitored even in crude bacterial lysates, as long as the absorbance peaks could be resolved. Using a similar strategy, it is possible to track DFHBI-1T by monitoring OD426 over time. In the case of DFHBI-1T, this is possible in the cultures themselves, as the peaks are resolvable even in intact cultures (data not shown). To quantify DFHBI-1T in cultures, OD426 measurements were normalised to those of the M9 media. Following this, the OD426 contribution of the bacterial cells were calculated using the OD700 measurements of cell number and conversions calculated from the absorbance spectra of cells (Csibra & Stan 2022). Subtracting the cellular OD426 contribution from the normalised OD426 allowed us to estimate the DFHBI-1T absorbance at 426 nm. This was converted to DFHBI-1T concentration via its extinction coefficient. The accuracy of this method is supported by the fact that starting concentrations of DFHBI-1T in all cultures were typically calculated within 10% of their intended concentrations (**Supplementary Fig. 2b**; which may have been due to imprecision in the exact mass of DFHBI-1T delivered or in pipetting, rather than in calculation).

### Ribosome structure visualisation

Molecular graphics and analysis was performed using ChimeraX (Meng et al. 2023). For all figures shown the PDB model 8B0X was used (Fromm et al. 2023). Molecular structures were displayed in cartoon backbone representation with ribosomal proteins coloured beige and rRNA coloured grey. Individual atoms, in between which Broccoli was inserted, were displayed as spheres and colour-coded. In case insertion sites were not modelled in the structure, the two closest residues were displayed instead.

### Data analysis

Data analysis was performed using R version 4.0.3 with general data handling packages of the tidyverse (Wickham et al. 2019). Plate reader data analysis was carried out using Parsley and FPCountR (Csibra 2021, 2023; Csibra & Stan 2022, 2023). Flow cytometry data analysis utilised FlopR (Fedorec 2023; Fedorec et al. 2020). Sequence alignment analysis made use of the SILVA rRNA database (Quast et al. 2013) and the DECIPHER package (Wright 2016).

## Data Availability

All plasmid sequences are provided as **Supplementary Data 1**.

## Acknowledgements

B.K. was supported by an ERASMUS+ grant. G.H.S. was supported by an EPSRC PhD Studentship. T.E.G. was supported by a Royal Society University Research Fellowship grants UF160357 and URF/R/221008, UKRI grant BB/W012448/1, a Turing Fellowship from The Alan Turing Institute under EPSRC grant EP/N510129/1. E.C. was supported by a Seeds for Success grant from the Imperial College Postdoc and Fellows Development Centre (2022) and an Eric Reid Fund for Methodology from the Biochemical Society (2023).

## Author Contributions

E.C. and T.E.G. conceived the project, provided guidance on experimental design and data analysis, supervised the work and should be considered joint senior authors. B.K., G.H.S. and E.C. performed experiments. E.C, G.H.S and B.K carried out data analysis. E.C. drafted the initial manuscript with input from all authors, and all authors contributed to the final manuscript.

