## Supplementary Figures for "Engineering orthogonal ribosomes for real-time monitoring using fluorescence"

### Table of Contents


| | Page |
|---|---|
| Supplementary Figure 1: Performance of mutant TO-ribosomes in two *E. coli* strains | 2 |
| Supplementary Figure 2: Tracking DFHBI-1T in bacterial cultures over time | 3 |
| Supplementary Figure 3: Internal initiation site in mCherry reduces signal to noise ratio of oRiboT2 | 4 |

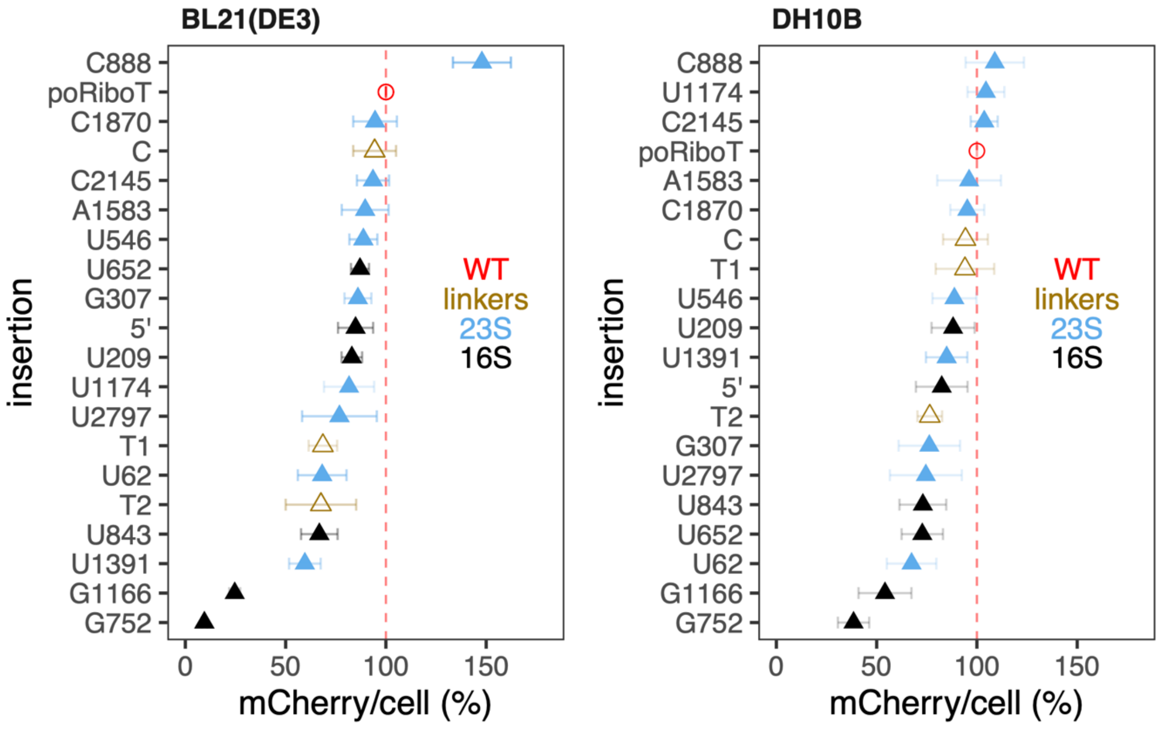


**Supplementary Figure 1: Performance of mutant TO-ribosomes in two *E. coli* strains.** Expanded data set from **Fig. 3a**, showing BL21(DE3) data (**a**) and DH10B data (**b**), with each insertion site ranked by performance. Data shows the mean and standard deviation of results over at least 2 independent experiments conducted in quadruplicate. Sites are coloured by ribosomal location: no insert/WT, red; insert in linker region, yellow; insert in 23S rRNA region, light blue; insert in 16S rRNA region, dark blue.

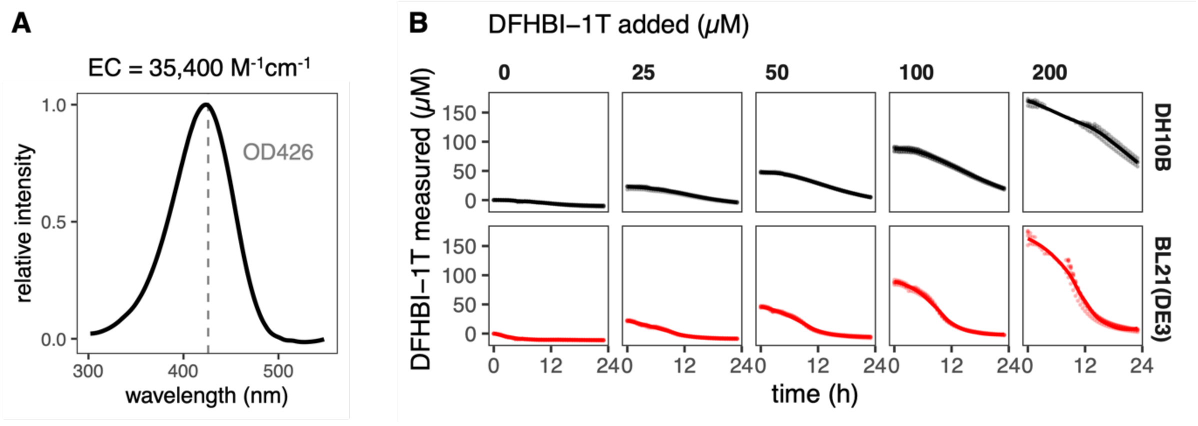


**Supplementary Figure 2. Tracking DFHBI-1T in bacterial cultures over time.** (**a**) Absorbance of DFHBI-1T (extracted from data in Filonov et al., 2015, Figure 6B). (**b**) DFHBI-1T levels measured over 24h in DH10B (black) and BL21(DE3) (red) cells containing no plasmid vectors. Data is presented as mean of triplicate wells (line) with individual triplicate data points (points), across a range of DFHBI-1T starting concentrations. EC, extinction coefficient.

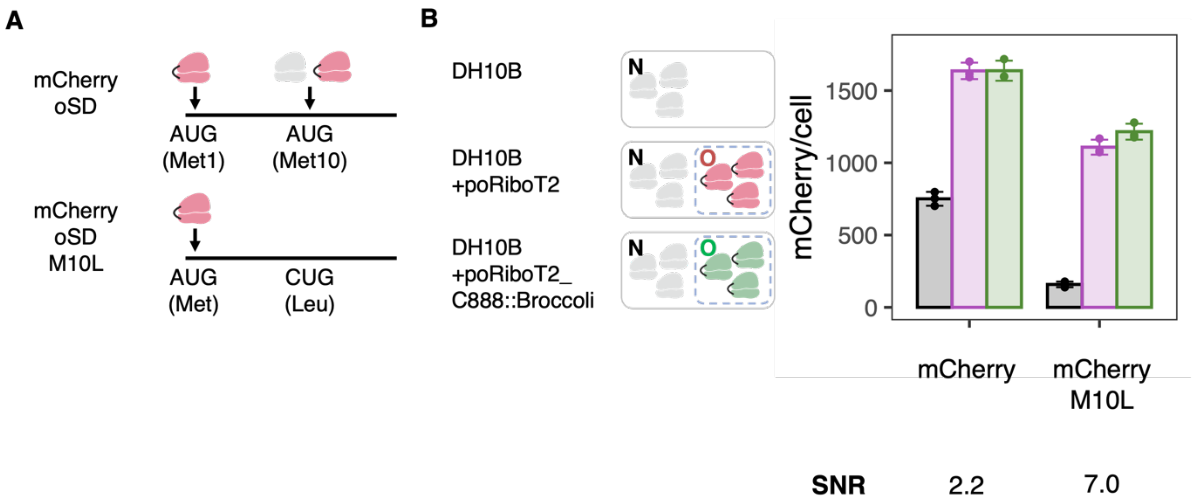


**Supplementary Figure 3. Internal initiation site in mCherry reduces signal to noise ratio of oRiboT2.** (**a**) Cartoon showing the canonical (blue) and putative internal (red, according to Fages-Lartaud et al., 2022) initiation sites, in the standard mCherry oSD construct (pS361_araoSDmCherry, top) versus the M10L variant (pS361_araoSDmCherryM10L, bottom). (**b**) mCherry abundance in cells without O-ribosomes (grey, DH10B) or with poRiboT2 (pink) or with poRiboT2 labelled with Broccoli (green). Quantifications were extracted from time course plate reader data from 720 min, present mean and standard deviations of triplicate wells, and are representative of two independent experiments. N, native (ribosomes); O, orthogonal (ribosomes); oSD, orthogonal Shine Dalgarno sequence; SNR, signal to noise ratio.
